# Tissue spatial correlation as cancer marker

**DOI:** 10.1101/340372

**Authors:** Masanori Takabayashi, Hassaan Majeed, Andre Kajdacsy-Balla, Gabriel Popescu

## Abstract

We propose a new intrinsic cancer marker in fixed tissue biopsy slides, which is based on the local spatial autocorrelation length obtained from quantitative phase images. The spatial autocorrelation length in a small region of the tissue phase image is sensitive to the nanoscale cellular morphological alterations and can hence inform on carcinogenesis. Therefore, this metric can potentially be used as an intrinsic cancer marker in histopathology. Typically, these correlation length maps are calculated by computing 2D Fourier transforms over image sub-regions – requiring long computational times. In this paper, we propose a more time efficient method of computing the correlation map and demonstrate its value for diagnosis of benign and malignant breast tissues. Our methodology is based on highly sensitive quantitative phase imaging data obtained by spatial light interference microscopy (SLIM).

## 1 Introduction

According to the World Health Organization (WHO), cancer is a major cause of death globally.^1^ Effective treatment strategies require early and accurate diagnosis of the disease. The gold standard method for cancer diagnosis is the microscopic investigation of a stained tissue biopsy by a trained clinical pathologist. Through this investigation, the pathologist looks for morphological signatures of either normal or abnormal tissue. Being qualitative, this type of assessment not only leads to inter-observer discrepancy but automation of some or part of the process through machine learning and image analysis is complicated by stain variability.^2, 3^ Ensuring consistency in the disease signatures extracted through image analysis of stained tissue remains a significant challenge due to variations in tissue preparation.^4^

Quantitative phase imaging (QPI)^5^ is a label-free microscopy technique where contrast is generated by the optical path-length difference (OPD), which is the product of the local thickness and refractive index of the specimen.^5–7^ For thin specimen, such as a tissue histology, the thickness can be considered spatially invariant, in which case QPI images are proportional to a mean refractive index map,^8, 9^ i.e., refractive index map integrated along z-axis. Since the refractive index is proportional to the dry mass content of cells and cellular matrix, it informs on tissue density as well as cell organization within tissue.^10, 11^ Tissue refractive index based markers have been used in the past for medical diagnosis and prognosis of several types of cancers and diseases.^12–24^ By generating contrast label-free, QPI lends itself much more readily to automated image analysis than bright-field microscopy, since stain variation is no longer an issue.^13^

In addition to the advantages of label-free imaging, the novel contrast mechanism in QPI provides access to additional, novel markers of disease, of value to histopathology.^14, 25^ In particular, since QPI systems employ interferometric measurements, they are sensitive to sub-wavelength fluctuations in OPD in both space and time.^5^ Therefore, local fluctuations in quantitative phase images inform on nanoscale morphological alterations of cell structures due to the dry mass accumulation as well a changes in extracellular matrix components. The tissue metric referred to as” disorder strength “, a marker of the spatial fluctuations of refractive index, i.e., nanoscale morphological alterations, was first used as a marker for pancreatic cancer diagnosis by Subramanian et al.^26^ Their group used a spectroscopic imaging modality to measure this marker and have subsequently employed it in diagnostic studies related to prostate, colon, breast, lung and other cancers.^27–36^ Thereafter, Eldridge et al. successfully extracted the disorder strength from quantitative phase images and demonstrated the relationship of the marker to cancer cell mechanical properties.^37^ They applied this analysis to colon, skin and lung cancer cells to demonstrate an inverse relationship between shear stiffness and disorder strength. Building on these results, A. Muoz et al. used QPI to study the on-set and progression of shear stiffness changes during malignant transformation in bronchial epithelial cells.^38^ Our group also showed that the disorder strength measured by spatial light interference microscopy (SLIM), a sensitive white light QPI method, is a quantitative marker of malignancy that can be used to classify benign and malignant breast tissue microarray (TMA) cores.^8^

In this paper, we propose the local spatial autocorrelation length as a new intrinsic marker of nanoscale morphological alteration in fixed tissue biopsies. Since the spatial autocorrelation length is related to the spatial refractive index fluctuations, which has been shown to detect malignancy in previous works, the length in a local region of tissue can be correlated with carcinogenesis. In the past, the local spatial autocorrelation length map was computed by calculating the 2D correlation function over regions of an image leading to long computation times. In this work, we present a more efficient algorithm for calculating the local spatial autocorrelation length map, requiring a smaller number of calculation steps. We then classify benign and malignant breast TMA cores using the local spatial autocorrelation length calculated by the proposed algorithm.

## 2 Materials and Methods

### 2.1 Spatial Light Interference Microscopy (SLIM)

The phase image, *ϕ*(*x*, *y*), measured in QPI is given by the expression

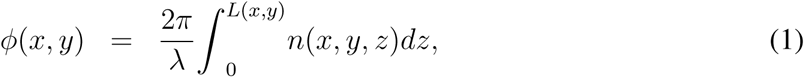

where *n*(*x, y, z*) is the refractive index contrast between the tissue and the surrounding medium, *L*(*x*, *y*) is the thickness of the tissue and *λ* the illumination wavelength. Here, we note that the intra-tissue refractive index fluctuations are integrated along *z* direction. In this work, we use 4 *µ*m thick tissue slices with small lateral variation in thickness (*L*(*x*, *y*) ≈ *L*) as the specimen.

A schematic of the SLIM setup is shown in Fig. 1(a). The SLIM module is attached to a commercial phase contrast microscope (PCM). The lamp filament is imaged onto the condenser annulus (Köhler illumination conditions) which is located at the front focal plane of the condenser lens. The specimen is located at the back focal plane of the condenser lens, and front focal plane of the objective. The scattered and unscattered fields are relayed by the objective and tube lenses. As a result, the expanded phase contrast image which has the intensity distribution in accordance with the phase contrast caused by the specimen is observed at the image plane. However, because the output of PCM is qualitative, the phase image, *ϕ*(*x*, *y*), cannot be directly retrieved from this image. The SLIM module extracts *ϕ*(*x*, *y*) by phase modulating the incident light with respect to the scattered light. The field at the image plane is Fourier transformed by the lens L1, such that the unscattered light can be spatially isolated from the scattered light. Since the incident light has the ring form, by displaying the corresponding ring pattern on the reflective liquid crystal phase modulator (LCPM), we ensure that the scattered light remains unaffected. Four phase shifts are applied to the unscattered light at increments of *π/*2 rad, as shown in Fig. 1(b). The corresponding four images captured by the charge coupled device (CCD) are obtained. Consequently, the quantitative phase image is retrieved as described in Ref. 5. Figure 1(c) shows the quantitative phase image and its expanded view of benign and malignant breast tissue samples.

**Fig 1.**
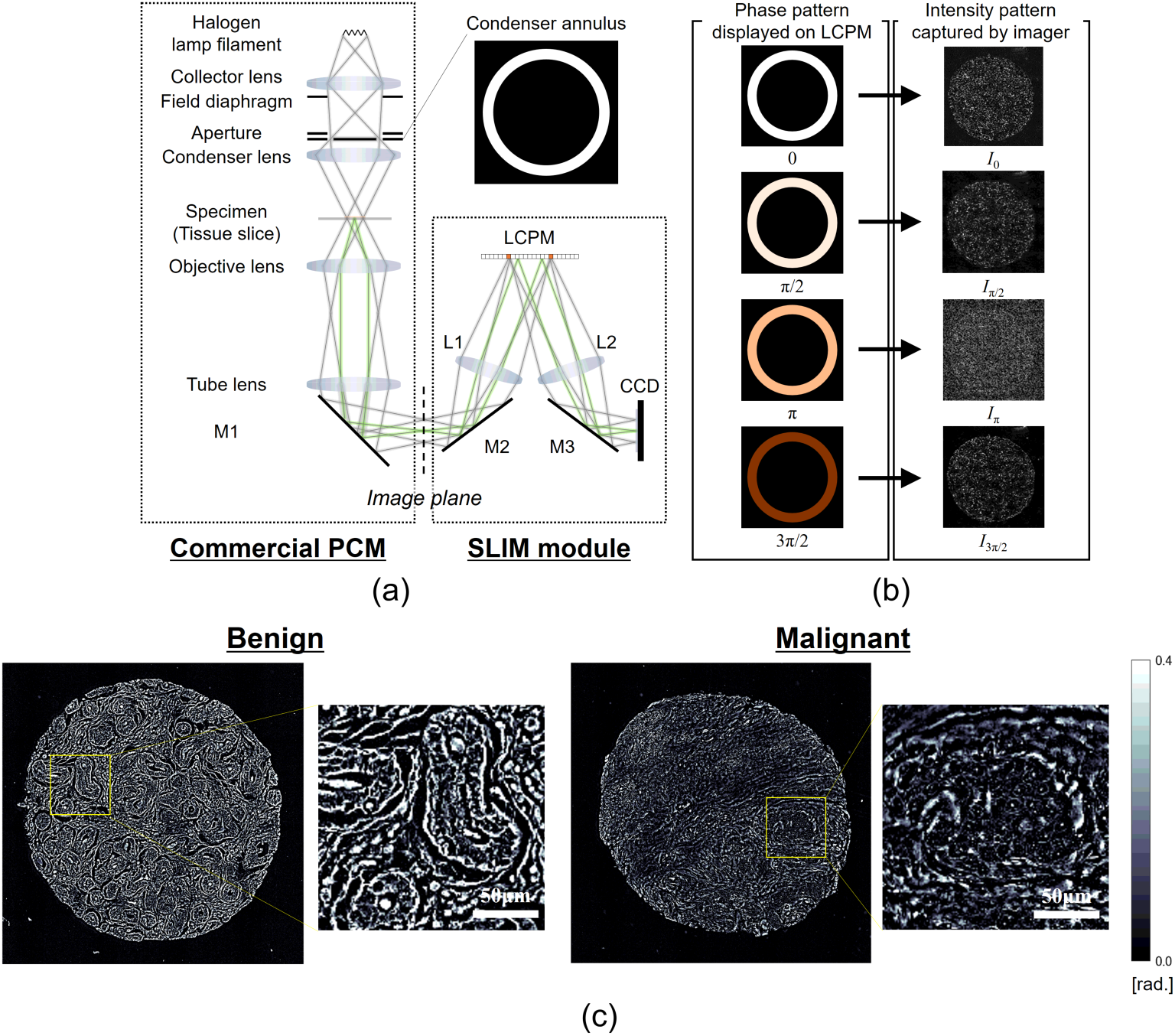
SLIM system. (a) Optical setup. (b) Phase patterns displayed on LCPM and corresponding intensity patterns captured by CCD. (c) Example of quantitative phase images of benign and malignant breast tissue cores and their expanded views.

### 2.2 Breast tissue microarrays

The samples comprised a TMA of cores constructed from breast tissue biopsies of 400 different patients. Each biopsy was formalin fixed and paraffin embedded before sectioning it into slices of 4 *µ*m thickness each using a microtome. Two parallel, adjacent sections were selected from each biopsy and one of these sections was stained using H&E, leaving the other one unstained. Cores were then constructed for both the stained and unstained tissue, and these were mounted on separate slides after de-paraffinization, using xylene as the mounting medium. The stained samples were imaged using a bright-field microscope, and their images were used by a board certified pathologist for diagnosing each core. In this paper, we studied benign tissue as well as cancerous tissue from 3 different grades: benign (*N* = 20), malignant (grade 1, *N* = 16), malignant (grade 2, *N* = 16) and malignant (grade 3, *N* = 14). Each patient consented to their tissue samples being used as a part of the study and the process of obtaining consent was approved by the Institute Review Board (IRB Protocol Number 2010-0519) at University of Illinois at Chicago (UIC). The data analysis was conducted on the samples at the University of Illinois at Urbana-Champaign (UIUC) after all patient identifiers had been removed. The procedures used in this study for conducting experiments using human subjects were also approved by the institute review board at UIUC (IRB Protocol Number 13900).

### 2.3 Formulation for local spatial autocorrelation length map

As mentioned, the local spatial autocorrelation length depends on the morphological disorder, i.e., local refractive index fluctuations. When the refractive index is spatially disordered, the spatial autocorrelation length within the local area will shorten. In general, the spatial autocorrelation length is calculated as the width of the spatial autocorrelation function. According to the Wiener-Khinchin theorem, the 2D spatial autocorrelation function can be obtained by taking inverse 2D Fourier transform of the spatial power spectrum. In other words, two 2D Fourier transforms for each image, leading to long computation times. Thus, to avoid this problem, we propose a new procedure that performs these calculation in the frequency-domain.

First, as shown in Fig. 2, we define the local spatial autocorrelation function as

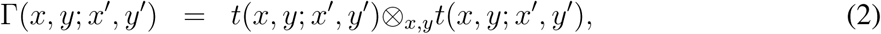

where ⊗_*x,y*_ denotes the 2D correlation operation over (*x*, *y*). Function *t*(*x*, *y*; *x*′, *y*′) is a local phase function centered at (*x*′, *y*′) and is expressed as

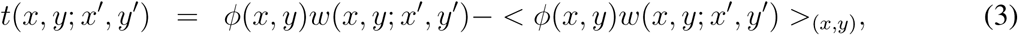

where 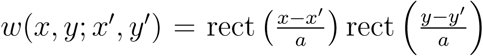 is a local window function centered at (*x*′, *y*′), of width of *a*. The angular brackets denote averaging within the local window.

**Fig 2.**
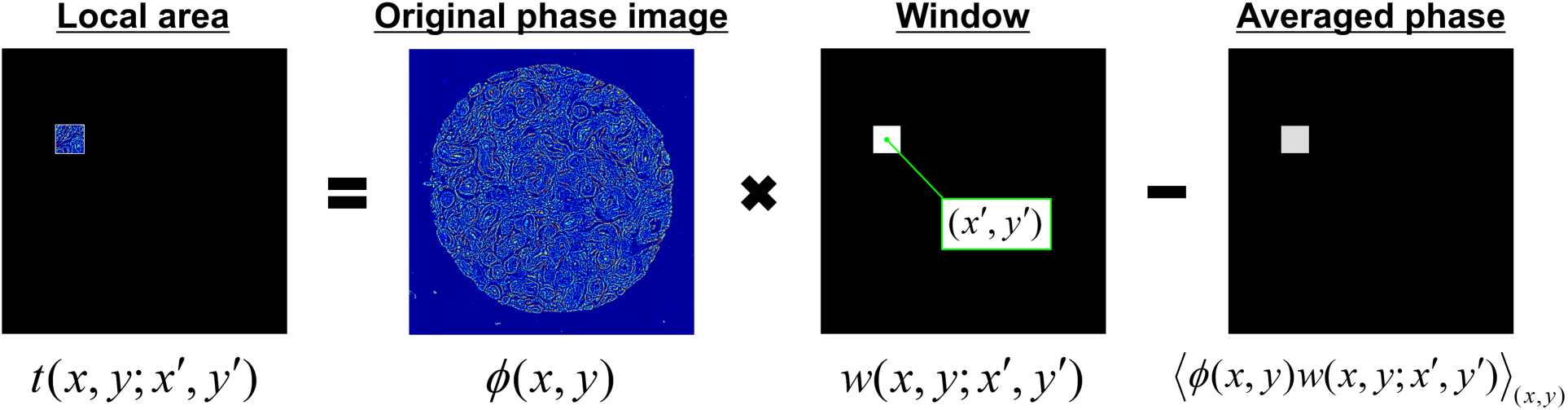
Definition of local 2D function *t*(*x*, *y*; *x*′, *y*′).

Next, we define the local spatial autocorrelation length map, *ρ*(*x*′, *y*′), as the variance of the probability density which can be obtained by normalizing Γ(*x*, *y*; *x*′, *y*′) by ∫∫ Γ(*x*, *y*; *x*′, *y*′)*dxdy*:

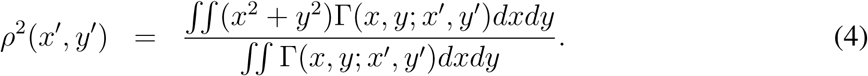

Here, *ρ*(*x*′, *y*′) can be related to the bandwidth map of the spatial power-spectrum, *τ* (*x*′, *y*′), as *ρ*(*x*′, *y*′)*τ* (*x*′, *y*′) = 2*π*. The local bandwidth, *τ* (*x*′, *y*′), itself is defined as

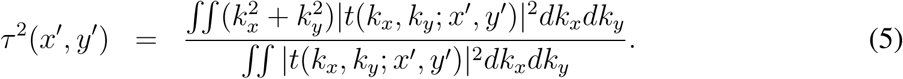

where *t*(*k*_*x*_, *k*_*y*_; *x*′, *y*′) is the Fourier transform of *t*(*x*, *y*; *x*′, *y*′) along (*x*, *y*). Using the differentiation property of Fourier transforms as well as Parseval ‘s theorem, this equation can be rewritten as

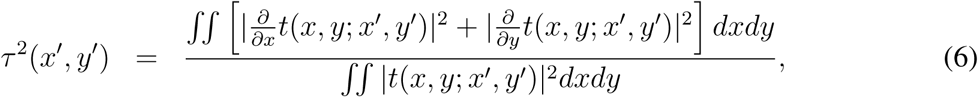

Finally, we can obtain the final result as

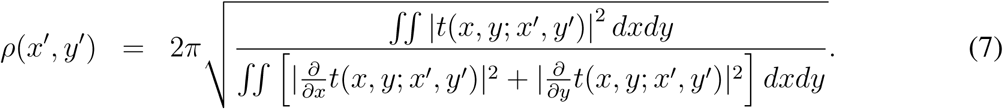

Using Eq. 7, the local spatial autocorrelation length maps can be calculated as shown in Fig. 3(b), which were obtained from the phase maps of benign and malignant cores with different grades (grade 1, 2 and 3) shown in Fig. 3(a). We used the local window with *a* = 64 pixels (8 *µ*m).

**Fig 3.**
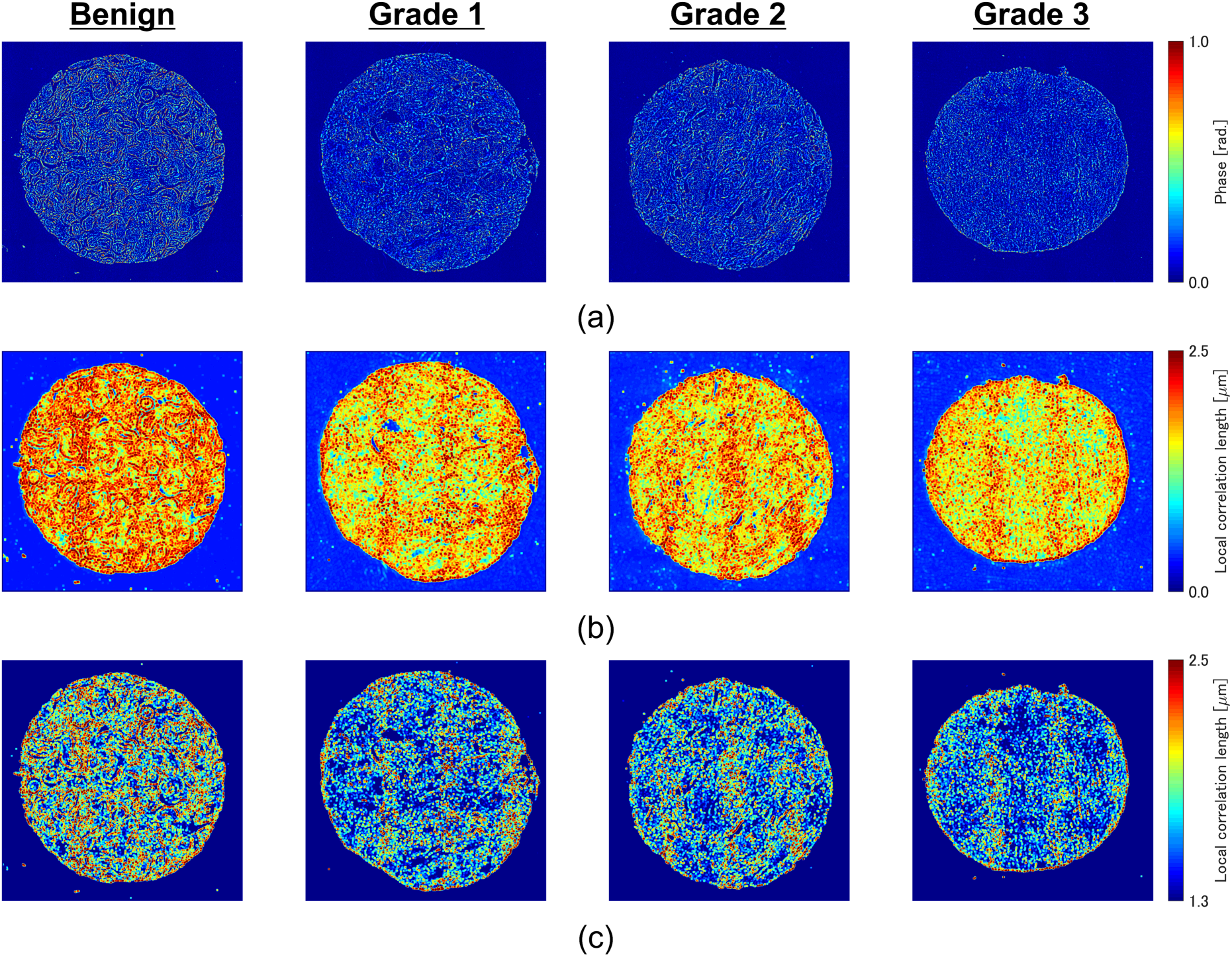
Example of local spatial autocorrelation length maps. (a) Quantitative phase images. (b) The local correlation length maps. (c) The local correlation length maps after applying the mask removing *ρ*(*x*′, *y*′) *<* 1.3*µ*m.

## 3 Results

We demonstrate that malignant transformation is correlated with metrics calculated from local correlation maps. The dataset we analysed consisted of 20 benign, 16 grade 1, 16 grade 2 and 14 grade 3 malignant cores. As the feature quantities, we use the value which is obtained by deviding the average by the standard deviation of the local correlation length. To extract only tissue regions, the background pixels were segmented out by setting a threshold in the *ρ*(*x*, *y*) map. This threshold value was determined empirically, and all pixels having correlation lengths below 1.3 *µ*m were treated as background. The maps obtained after the background pixels reduction process are shown in Fig. 3(c). This calculation took approximately 45 min. per one core which consists of 7040 *×* 7040 pixels (880 *×* 880*µ*m^2^) using PC with Intel Core i5-3470 CPU (3.20 GHz), 16.0 GB RAM whereas it was estimated to take 90 min. without our algorithm, i.e., with the algorithm using fast Fourier transform on the basis of Wiener-Khinchin theorem. This calculation time can be improved, for example by using GPU acceleration.

Figure 4 compares the average divided by the standard deviation of *ρ* map between benign and malignant cores (grade 1, 2 and 3). The p-values which were obtained by two-sided Wilcoxon ranksum test are listed in Table 1. Since the p-value between 20 benign and 46 malignant cores was 0.000876, the local correlation length correlates with cancer grades. Furthermore, the results indicate that the statistically significant differences between cores with more than 2 inter-grade differences. This means that this label-free marker can be used for separating the lower-risk cases (benign and grade 1) from the higher-risk cases (grade 2 and 3). However, it may need to be combined with other markers for more detail separation of grades. We conclude that the local correlation map can potentially be used by clinical pathologists as a supplementary label-free disease marker for gauging the onset of malignancy especially in borderline cases.

**Table 1.** The p-values between different grades

| Grades to be evaluated | p-value |
| --- | --- |
| Benign - Malignant (G1, G2, G3) | 0.000876 |
| Benign and Malignant (G1) - Malignant (G2, G3) | 0.000335 |
| Benign - Malignant (G1) | 0.101101 |
| Benign - Malignant (G2) | 0.005891 |
| Benign - Malignant (G3) | 0.000498 |
| Malignant (G1) - Malignant (G2) | 0.193509 |
| Malignant (G1) - Malignant (G3) | 0.018837 |
| Malignant (G2) - Malignant (G3) | 0.417581 |

**Fig 4.**
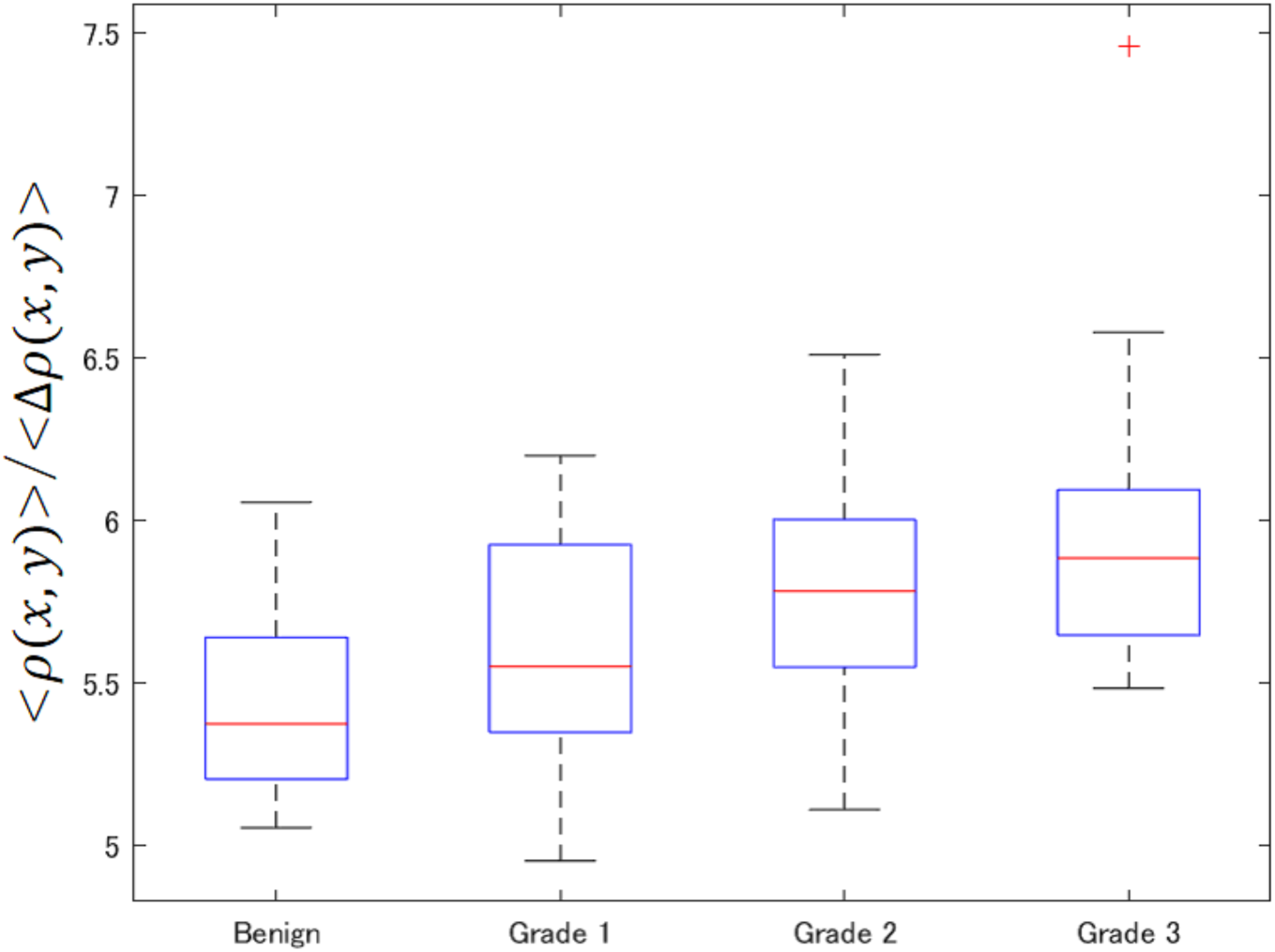
Local spatial autocorrelation length of benign (*N* =20) and grade 1 (*N* =16), grade 2 (*N* =16) and grade 3 (*N* =14) tissues.

## 4 Summary and conclusion

In summary, we have presented an efficient algorithm for the computation of the local correlation length within refractive index maps of fixed breast tissue biopsy slides. This length metric describes the local refractive index fluctuations within the tissue specimen. Since in this work the refractive index maps are extracted using SLIM, which has sub-nanometer optical path length sensitivity, the correlation length is indicative of nanoscale cellular morphology. Standard computation of correlation length maps involves 2D Fourier transforms which can lead to long computation times, especially for large analysis window sizes. We improve calculation throughput by performing part of the computation in the frequency domain.

A comparison of the extracted correlation lengths between benign and malignant TMA cores showed that this metric is on average smaller for malignant cores indicating increased randomization of tissue morphology (as captured by its refractive index). Statistically significant differences in correlation lengths were observed between the two classes (*N* =20 for benign and *N* =46 for malignant) indicating that this label-free disease marker can potentially be used by clinical pathologists for gauging the onset of malignancy especially in borderline cases. Furthermore, although the separation between neighboring grades cannot be achieved without the help of other markers, the results indicated that the local correlation length can also be used for separating the lower-risk cases (benign and grade 1) from the higher-risk cases (grade 2 and 3).

On the other hand, there is room for improvement on the calculation time and the screening accuracy. For example, calculation acceleration by GPU will drastically improve the calculation time and the optimization of local window size and combining with other markers such as disorder strength^8^ will contribute to improve the screening accuracy. Because SLIM can be implemented as an upgrade of the existing microscopes, the extraction of intrinsic markers from quantitative phase images obtained through SLIM is expect to be plugged into the existing pathology work flow. Tissue spatial correlation information can add to the existing toolbox that the pathologists already have and help improve diagnosis accuracy and objectivity.

## Acknowledgments

We would like to thank Prof. Garth H. Rauscher for his pathology expertise. This work was supported by National Science Foundation (CBET-0939511 STC, DBI 1450962 EAGER, IIP-1353368, and CBET-1040461 MRI), and by JSPS KAKENHI Grant Number 18K14150.

This work is based partly on scientific content previously reported in SPIE proceedings: Masanori Takabayashi, Hassaan Majeed, Andre Kajdacsy-Balla, Gabriel Popescu, “High throughput calculation of local spatial autocorrelation length for label-free diagnosis of tissue biopsy”, Proc. SPIE 10503, Quantitative Phase Imaging IV, 105032B; doi: 10.1117/12.2293991;

**Masanori Takabayashi** is an associate professor at Kyushu Institute of Technology. He received his BE, ME and PhD degree in information science and technology from Hokkaido University in 2007, 2009 and 2012, respectively. His current research interests include volume holography, digital holography, and quantitative phase imaging for medical diagnosis.

